# Polygenic Risk Associated with Alzheimer’s Disease and Other Traits Influences Genes Involved in T Cell Signaling and Activation

**DOI:** 10.1101/2023.05.10.540219

**Authors:** Dallin Dressman, Shinya Tasaki, Lei Yu, Julie Schneider, David A. Bennett, Wassim Elyaman, Badri Vardarajan

**Affiliations:** Department of Neurology, Columbia University, New York, NY; The Taub Institute for Research on Alzheimer’s Disease and the Aging Brain, Columbia University, New York, NY; Rush University Medical Center, Rush Alzheimer’s Disease Center, Chicago, IL; Rush University Medical Center, Department of Neurological Sciences, Chicago, IL; Rush University Medical Center, Department of Pathology, Chicago, IL; College of Physicians and Surgeons, Columbia University, The New York Presbyterian Hospital, The Gertrude H. Sergievsky Center, New York, NY

## Abstract

T cells, members of the adaptive immune system known for their ability to respond to an enormous variety of pathogens and other insults, are increasingly recognized as important mediators of pathology in neurodegeneration and other diseases. Previously, we and others have shown that T cell gene expression phenotypes are regulated by genetic variants associated with autoimmune disease, neurodegenerative disease, and inflammatory processes. However, many complex diseases have polygenic risk with thousands of common variants contributing a small amount to disease heritability. Here, we compute the polygenic risk score (PRS) of several autoimmune, neurological, and psychiatric disorders and present the first correlation of these PRSs with T cell gene expression, using transcriptomic and genomic sequencing data from a cohort of Alzheimer’s disease (AD) patients and age-matched controls. We validate our AD PRS against clinical metrics in our cohort and then compare PRS-associated genes across traits and four T cell subtypes. Several genes and biological pathways associated with the PRS for these traits relate to functions such as T cell chemotaxis, differentiation, response to and production of cytokines, and regulation of T cell receptor signaling. We also found that the trait-associated gene expression signature for certain traits was polarized towards a particular T cell subset, such as CD4+ for autoimmune disease traits or CD8+ for some psychiatric disease traits. Our findings may help guide efforts in precision medicine to target specific T cell functions in individuals with high polygenic risk for various complex diseases.

## Introduction

Alzheimer’s disease (AD) is a chronic neurodegenerative condition that afflicts millions of Americans. While AD pathology has long been known to include aberrant aggregation of amyloid beta peptide and tau protein, it is also linked to inflammation and other immune processes. T cells, which form part of the adaptive immune response to pathogens and other biological insults, have displayed several key phenotypic changes in recent studies using bio-samples from human AD patients. Specifically, T cells in AD patients show a high degree of clonal expansion and a less diverse T cell receptor repertoire, increased infiltration into the cerebrospinal fluid (CSF) and brain parenchyma, and upregulation of genes involved in cytotoxicity, inflammation, immunosenescence, and response to certain chemokines (1–3). We previously detailed expression quantitative trait loci (eQTL), or T cell gene expression changes correlated with single nucleotide polymorphisms, in AD patients and age-matched controls (4). We identified numerous cis and trans-eQTL distributed across the genome, showing that many variants far from a gene locus can regulate gene expression. Aggregating the effects of many genetic variants regulating gene expression may better capture genotype-phenotype correlation in complex disease, especially if these variants have been previously linked to disease risk. Thus, we now focus on correlating polygenic risk scores (PRSs) for AD and other conditions with our T cell gene expression data, building on our previous findings and on research using polygenic risk scores to better understand AD and related phenotypes.

When genetic and AD diagnosis data is available for a cohort, the predictive value of a PRS for AD can be calculated based on clinical diagnosis (determined by cognitive examination) or pathological diagnosis (determined by a threshold of pathological biomarkers, such as plaques or tangles, in post-mortem brain samples and Aβ40/42 or pTau181 levels in CSF of plasma antemortem). Depending on the study and the choice of diagnostic strategy, PRS predictive value for AD in previous literature ranges from 72-84% (5–9). These and other studies also consistently report a higher incidence of AD among individuals with high PRS (10,11), validating the correlation of the AD PRS with disease status. At present, PRS alone is insufficient for prediction of AD in a clinical setting. However, several groups are testing approaches to achieve higher accuracy with combined models using PRS alongside other metrics such as AD biomarkers or cognitive performance.

Polygenic risk for AD has also been correlated with quantitative measures of cognitive performance and AD neuropathology. Several groups have enrolled cohorts of non-aged individuals and followed them longitudinally, showing that an AD PRS can help predict the conversion of mild cognitive impairment to AD (12–14). Other studies have shown that cognitive performance in non-AD individuals is still worse in individuals with high PRS for AD (10,15,16), even in children and adolescents (17). Individuals with high polygenic risk for AD are also more likely to display biomarkers of advanced AD neuropathology, including higher amyloid signal on positron emission tomography (15,18), reduced volume in hippocampus, amygdala, and entorhinal cortex on magnetic resonance imaging (10,11,15,17,18), increased levels of amyloid beta and tau in plasma or cerebrospinal fluid (10,19–21), and increased activity of gamma-secretase (19), an enzyme that helps process APP into Aβ42. Thus, individuals with high polygenic risk for AD, even before showing the onset of clinical dementia, show signs of AD progression and reduced cognitive capacity.

While novel approaches to PRS studies are accelerating, few studies have correlated PRSs with gene expression at this stage (22–24), and no studies, to our knowledge, have correlated a PRS for any disease with T cell gene expression. We hypothesize that disease relevant genes and pathways will be differentially expressed at varying levels of polygenic risk of the disease. In contrast to traditional QTL approaches, which aim to identify genetic variants that regulate gene expression in a phenotype agnostic manner, association of PRS with gene expression will identify differential expression by overall genetic risk of disease. Thus, we calculate a PRS for AD, validating it against diagnostic data and neuropathological measurements. We then calculate a PRS for 18 other traits related to immune function, autoimmune disease, neurological disease, and psychiatric disorders. We correlate these PRSs with RNA-sequencing data from four T cell subsets sorted from peripheral blood samples from AD patients and age-matched controls in the Religious Orders Study and Rush Memory and Aging Project (25).

We aim to understand the differential gene expression in T cells at different polygenic risk levels for AD and other disorders. We expect that our dataset, involving four T cell subsets, will highlight differences in PRS-associated genes across T cell subtypes and disease traits. Our findings highlight biological pathways and other mechanisms of polygenic risk for disease, which can aid in hypothesis generation for future targeted studies of T cell behavior in AD and other conditions. They also provide interesting comparisons to previous genotype-phenotype correlation studies using T cell RNA-sequencing data, from our group and others (4,26–32).

## Results

### Calculation and validation of a polygenic risk score for AD

We first used PRSice-2 (33) to calculate a genome-wide polygenic risk score for AD, for 2051 individuals in ROSMAP. We used the summary statistics from the Kunkle et al. 2019 (34) genome-wide association study (GWAS) for SNP effect sizes excluding SNPs in the APOE locus (see Methods). PRSs were calculated using SNPs associated with AD at *p*-value thresholds ranging from 5 x 10^-8^ to 1 (all SNPs), and PRS distributions at each threshold were standardized to have a mean of 0 and standard deviation of 1. For individuals with clinical or pathological AD diagnostic data, we computed the predictive value of the AD PRS for each p-value threshold (see **Supplementary figure 1**). The maximum predictive value of the AD PRS was 78.19% for clinical AD, and 69.55% for pathological AD. These values are likely inflated due to the overlap between the 2051 ROSMAP individuals and subjects in the Kunkle et al. 2019 GWAS used as base data. Calculation of the AD PRS using only non-overlapping subjects yielded a predictive value of 53.59% for clinical AD.

For additional validation of the utility of the AD PRS, we correlated the AD PRS with neuropathological traits. The AD PRS was significantly associated with Braak staging (35), a semi-quantitative measure of the spread of tau pathology, using one-way ANOVA (Kruskal-Wallis *p* < 0.0001). Linear regressions of the AD PRS against quantitative measures of amyloid burden, tau tangles, and global pathology (gpath) using age, sex, and the first ten principal components from genotyping data as covariates, all had slopes significantly different from zero **(Supplementary figure 1).** From these analyses, we concluded that our AD PRS had reasonable predictive value for AD status and correlation with measures of AD pathology. We also found that using *p*-value thresholds of 0.75 or 1 for inclusion of SNPs in PRS calculation resulted in the highest predictive value for AD.

### Correlation of gene expression with PRSs for AD and other traits

Owing to our previous findings detecting eQTL with disease-associated variants in several trait categories (4), we computed PRSs for common traits or diseases in each of these categories in our dataset. Specifically, we calculated PRSs for lymphocyte counts (36), white blood cell counts (36), C-reactive protein levels (36), ulcerative colitis (37), Crohn’s disease (37), multiple sclerosis (38), rheumatoid arthritis (39), systemic lupus erythematosus (40), type 1 diabetes (41), Parkinson’s disease (42), amyotrophic lateral sclerosis (43), epilepsy (44), stroke (45), bipolar disorder (46), anxiety disorder (47), major depressive disorder (48), schizophrenia (49), and post-traumatic stress disorder (50).

78 of our 96 RNA-sequencing subjects had genotyping array data available for PRS calculation. One of these subjects was excluded because their AD PRS was an outlier at the PRS distribution for *p* = 1, leaving 77 subjects for correlation between PRSs and gene expression. Each subject had bulk RNA-sequencing data from the following T cell subsets, sorted using flow cytometry: CD4+CD45RO-, CD4+CD45RO+, CD8+CD45RO-, and CD8+CD45RO+. In these subjects, we computed the PRSs using all independent SNPs in the genome (*p*-value threshold of 1), since we did not have phenotypic data for PRS optimization, and the optimal PRS for AD was derived from the higher *p*-value thresholds. We then correlated the PRSs with T cell gene expression, using linear regression with age, sex, AD status, and the first 10 principal components from genotyping as covariates. We included only genes with expression in at least 20% of subjects, with a maximum expression count of at least 3, the same parameters used in our previous eQTL study (4). 46 of our sequencing subjects also have brain RNA-sequencing data from a recent publication (23), although we focus only on T cell gene expression here. A listing of all PRS-associated genes by trait and T cell subsets is found in **Supplementary Table 1**.

Genes with a nominally significant relationship with the PRS are shown on the heatmap in **Figure 1**. Overall, 6110 genes (of 6139 genes passing minimum expression thresholds) were associated with the PRS for at least one trait in at least one cell type. About 50% of these genes were significantly associated with the PRS in five or more trait/cell type combinations (see **Supplementary Figure 2** for breakdown of trait overlap across cell types and within cell types). Interestingly, several autoimmune disease and psychiatric disorder traits feature almost all significant PRS-associated genes in an inverse relationship with the PRS (as seen by cells colored blue), regardless of the cell type, suggesting a pattern of downregulated genes in individuals with high polygenic risk for these conditions.

**Figure 1.**
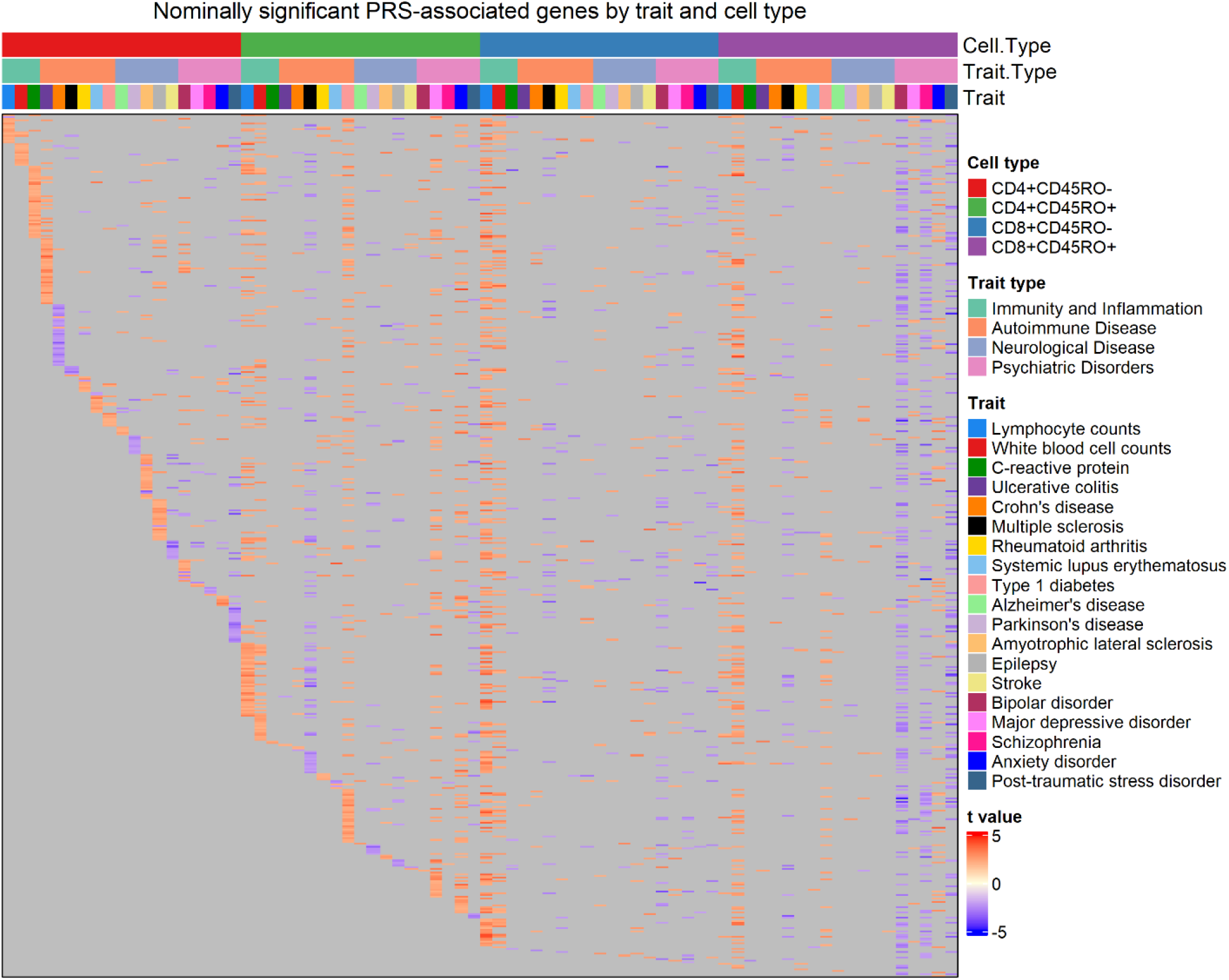
Heatmap of genes nominally associated with the PRS for one or more traits. Annotation rows above the heatmap show cell type, trait type, and trait, matching the color legends at the right. Each row of the heatmap is a gene, and each cell is colored by the strength and direction of the association with the PRS (shown by the color legend at bottom right), or gray if the association is insignificant.

Adjusting for multiple testing with the Benjamini-Hochberg false discovery rate method yielded 2462 significant genes at the q = 0.1 threshold, and 1217 at the q = 0.05 threshold, almost all of which resulted from the association of the PRS for lymphocyte counts with gene expression in CD8+CD45RO-T cells. Only two of these genes remained significant after more stringent Bonferroni multiple testing correction with q = 0.1: CRIP1, a protein involved in zinc transport, and SKIDA1.

364 genes were nominally associated with the PRS for over ten traits across T cell subsets, including genes that play a role in T cell function or disease pathology. TRAT1 helps stabilize the CD3/T cell receptor (TCR) signaling complex (51), while SLA negatively regulates TCR signaling (52). CD44 and CD69 are markers of T cell memory and activation, respectively (53,54), and MAF is associated with differentiation of CD4+ T cells into the interleukin (IL)-4 producing Th2 subtype (55). SOD1, a gene with risk variants for ALS (56), was associated with the PRS for lymphocyte counts, white blood cell counts, MS, type 1 diabetes, bipolar, and PTSD in one or more T cell subsets in our data. Finally, CXCR4, a chemokine receptor, was found to be upregulated in a cluster of T cells in the cerebrospinal fluid of Lewy body dementia patients (3) and is thought to regulate infiltration of T cells into the CNS in disease and aging (57,58). Interestingly, CXCR4 was not associated with the PRS for AD or PD in any of the T cell subtypes in our data.

We further compared PRS-associated genes across traits by calculating the Pearson’s correlation coefficient of the effect size of PRS association for any two traits within each T cell subset. As expected, genes associated with the PRSs for lymphocyte and white blood cell counts were highly correlated in all four T cell subtypes (see **Figure 2**, quantification in **Supplementary Table 2**). Within a particular trait category (immune function, autoimmune diseases, neurological conditions, and psychiatric disorders), significant correlations between trait-associated gene expression patterns were generally positive. Trends in correlation coefficients also remained the same across T cell subtypes, with few exceptions. However, the gene expression signature associated with the PRS for stroke was negatively correlated with other neurological conditions such as Parkinson’s disease, amyotrophic lateral sclerosis, and epilepsy. Interestingly, it bore a strong positive correlation to the PRS-associated genes for the psychiatric disorders we included, except schizophrenia.

**Figure 2.**
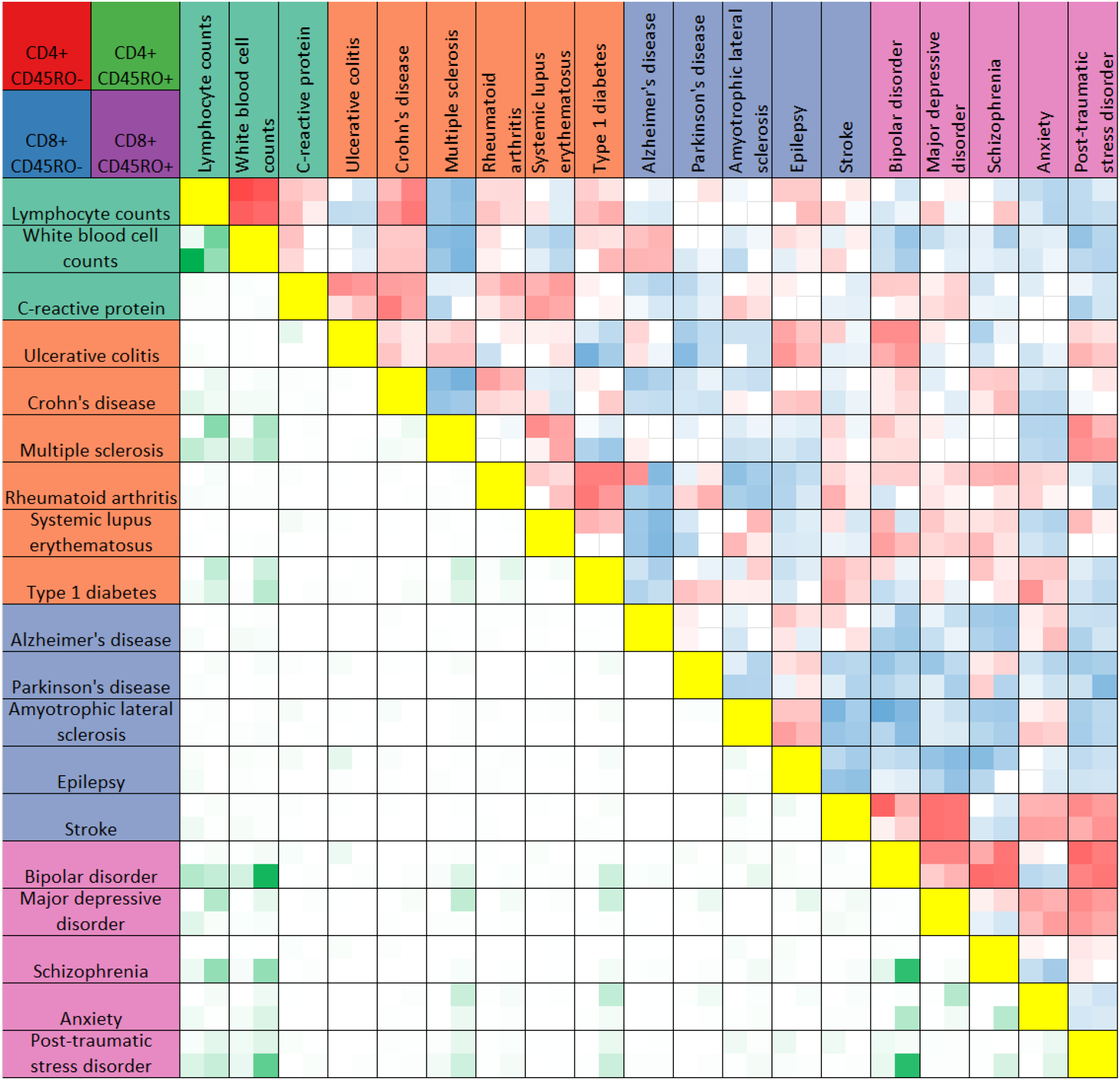
Heatmap comparison of PRS-associated genes across traits. The Heatmap summarizes results of Pearson’s correlation test between PRS-associated genes for any two traits (boxes above yellow diagonal) or the numbers of genes significantly associated with the PRS (*p* < 0.05) shared between any two traits (boxes below the yellow diagonal). Traits are listed above and to the left of the heatmap, colored according to trait categories as in Figure 2. Each box in the heatmap reflects four values, for CD4+CD45RO-(top left), CD4+CD45RO+ (top right), CD8+CD45RO-(bottom left) and CD8+CD45RO+ (bottom right). Boxes for Pearson’s correlation tests are darker red for r values approaching 1, darker blue for r values approaching -1, and white if insignificant after Bonferroni multiple testing correction with n = 171. Boxes comparing overlap of significant PRS-associated genes are darker green for higher numbers of shared genes between two traits. For quantification, see Supplementary Table 2.

### Pathway analysis of PRS-associated genes

To interrogate the functional connections of PRS-associated genes, we used Gene Set Enrichment Analysis (59) (GSEA) to detect biological pathways over-represented by genes from our dataset. The Gene Ontology Biological Processes database (60) is a curated list of genes involved in various biological functions, most of which are not specific to a particular cell type. We ran GSEA by supplying a list of genes, from each combination of T cell subset and disease trait, ranked by the *t* value for association with the PRS. GSEA detected pathways from GO Biological Processes over-represented by genes at the extreme ends of the *t* value distribution for each trait and T cell subset. Pathways with a false discovery rate q-value under 0.05 are shown in the four dot plots in **Figure 3** organized by cell type and colored within each plot by trait.

**Figure 3.**
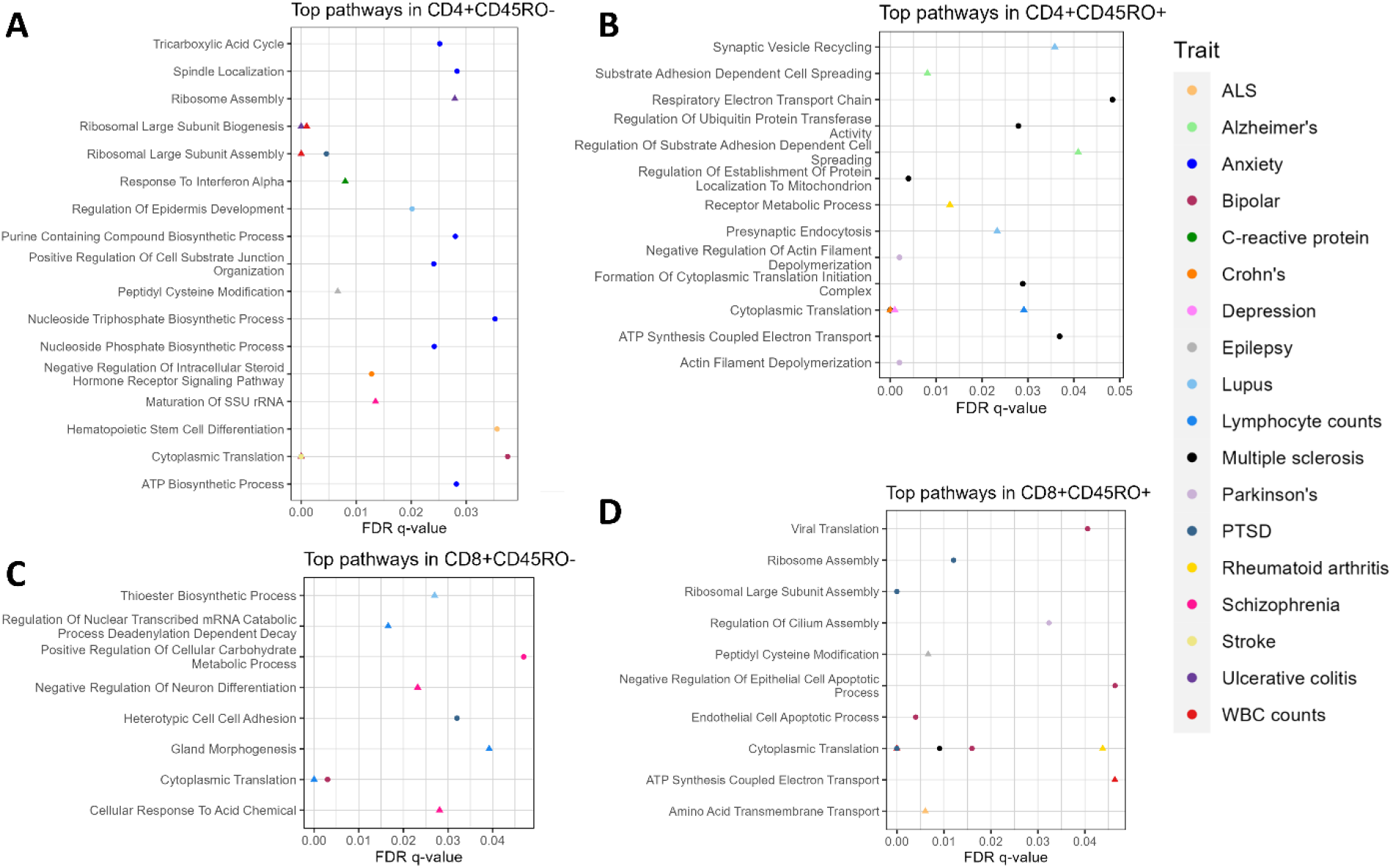
(previous page). GSEA of PRS-associated genes by T cell subset. Dot plots show pathways significantly over-represented after multiple testing correction for **A)** CD4+CD45RO-, **B)** CD4+CD45RO+, **C)** CD8+CD45RO-, and **D)** CD8+CD45RO+. Trait is shown by color (see legend at right), q-value after multiple testing correction is shown by position on the x-axis, and shape denotes whether the pathway is constructed of genes with a negative *t* value (circle) or positive (triangle). Statistics of significant GSEA pathways are found in supplementary table 3.

The majority of pathways significant after multiple testing correction relate to translation, RNA processing, cellular respiration, and oxidative phosphorylation, none of which is unique to T cell function (significant GSEA pathways can also be viewed in **Supplementary Table 3**). This is consistent with our pathway analysis findings for trans-eQTL genes (4), which also mainly returned basic cellular functions nonspecific to any one cell type. However, several pathways raised the possibility for interesting functional connections. In CD4+CD45RO-T cells, the “response to interferon alpha” pathway was associated with the PRS for C-reactive protein levels, and the “hematopoietic stem cell differentiation pathway” was associated with the PRS for ALS. Other PRS traits with notable pathway analysis findings included “negative regulation of neuron differentiation” and “cellular response to acid chemical” for schizophrenia in CD8+CD45RO-, “heterotypic cell-cell adhesion” for PTSD in CD8+CD45RO-, and “viral translation” for bipolar in CD8+CD45RO+.

Closer interrogation of these pathways revealed several genes involved in T cell and other immune cell functions. For “substrate adhesion-dependent cell spreading”, associated with the PRS for AD in CD4+CD45RO+, genes included C1QBP, a receptor that binds a protein in the complement system (61), and ITGA4, which regulates T cell homing and may affect T cell migration into the brain (62). Genes in the “cellular response to acid chemical” pathway include ZEB1, which inhibits translation of the IL-2 cytokine that is vital for T cells (63), KLF2, a transcription factor that prevents migration of naïve T cells into nonlymphoid tissues (64), MTOR, a key regulator of cell growth and proliferation (65), and several members of the LAMTOR gene family, which are among upstream regulators of the mTOR complex (66). The “heterotypic cell-cell adhesion pathway” includes several genes related to integrin signaling in T cells, which is key to their ability to extravasate from the circulation (67). It also involves genes that can modulate or serve as a marker of T cell activation and memory, such as PTPRC, CD44, and CD47 (68,69).

These findings suggest a potential dysfunction of T cells in disorders whose connection to T cell biology is sparsely studied, such as bipolar disorder or PTSD. Pathway analysis also sheds light on potential T cell mechanisms for traits whose connection to adaptive immunity is well known, such as lymphocyte counts or C-reactive protein levels. Finally, these data imply that the extent and nature of T cell activity in disease pathology may change with respect to polygenic risk for these conditions.

### Comparing abundance of PRS-associated genes between CD4+ and CD8+ T cell subsets

Some conditions feature pathological mechanisms that are unique to a particular T cell subset, or favored by one T cell subset over another. Such is the case for several disease traits in our PRS analysis. Many autoimmune disease traits, for example, are driven more by CD4+ T cell-mediated pathology than CD8+ (70), while recent single-cell sequencing research in neurodegenerative disease patients has highlighted mechanisms of inflammation and cytotoxicity in CD8+ T cell clusters (2,3,71). We sought to determine whether similar patterns existed in our genotype-phenotype correlation studies, based on the abundance of PRS-associated genes in CD4+ and CD8+ T cell subtypes by trait.

**Figure 4** shows that autoimmune disease traits (ulcerative colitis through type 1 diabetes) have most PRS-associated genes in CD4+ T cell subtypes. However, when only considering genes inversely related to the PRS (denoted by a negative t-value and shown in the rightmost column), several traits have the majority of these genes in CD8+ T cell subtypes. Traits with a higher percentage of all PRS-associated genes in CD8+ T cell subtypes include bipolar disorder, schizophrenia, post-traumatic stress disorder, lymphocyte counts, and white blood cell counts. For several traits, the relative percentage of genes in CD4+ or CD8+ T cell subsets differs widely by the *t*-value sign. For Parkinson’s, Crohn’s disease, and stroke, genes proportionally associated with the PRS are mainly in CD8+ T cells, while genes inversely associated with the PRS are predominantly in CD4+ T cells. The opposite is true for C-reactive protein levels, bipolar disorder, and schizophrenia (for quantification, see **Supplementary Table 4**).

**Figure 4.**
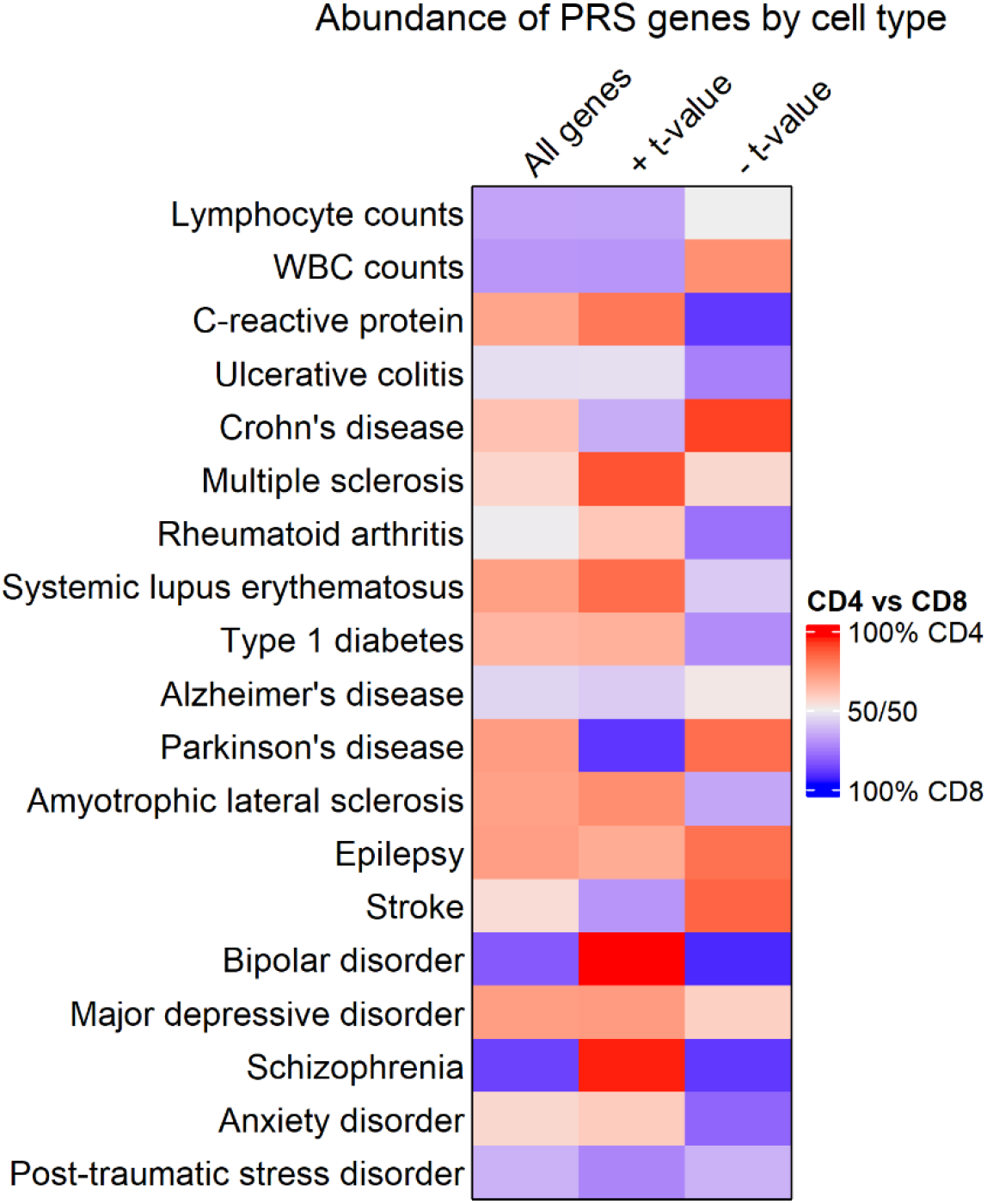
(previous page). Comparing enrichment of PRS-associated genes between CD4+ and CD8+ T cell subsets. Heatmap showing the relative abundance of all PRS-associated genes (left column), genes with a positive *t*-value (middle column), or genes with a negative t-value (right column) in CD4+ or CD8+ T cell subsets. Cells are colored red when most genes are in CD4+ T cell subsets, and blue when most genes are in CD8+. Quantities associated with this heatmap are available in supplementary table 4.

We looked further into PRS-associated genes on the extreme ends of the t value distribution for traits where gene enrichment in CD4+ vs. CD8+ T cells differed by *t* value sign. Genes in CD4+ T cells with a negative *t* value for association with the PRS included HAVCR1 in Crohn’s disease, a gene which mediates T cell trafficking in autoimmunity and inflammation (72). CD4+ genes inversely associated with the PRS for Parkinson’s included DOCK8, which assists in T cell chemotaxis (73), and CD59, a protein upregulated on activated CD4+ T cells (74). Among CD8+ genes that scaled proportionally with the PRS for Parkinson’s disease, top genes included GZMM, involved in CD8+ T cell cytotoxicity (75) and IL2RB, part of the IL-2 receptor complex.

Several genes related to the PRS for C-reactive protein levels often serve as regulators or downstream mediators of cytokine-induced signaling. These include IL16, IL10RA (a part of the IL-10 receptor complex), CASP1 (which cleaves the precursor of IL-1β, (76)), and PPM1A, which helps terminate TGFβ signaling and TNFα-induced NFκB signaling (77,78). PPM1A, while upregulated in CD4+ T cells of individuals with high PRS for C-reactive protein levels, is downregulated in CD8+ T cells for the same individuals in our data. Individuals with high polygenic risk for schizophrenia increased expression in CD4+ T cells of MAL, a gene downregulated in memory T cells (79), and STAM, a gene mediating downstream effects of cytokine signaling (80). Meanwhile, CD8+ genes inversely related with the PRS for schizophrenia included IFNAR1, part of the receptor complex that responds to type 1 interferons. Finally, in bipolar disorder, CD8+ T cells in individuals with a high PRS reduced expression of PTPRJ, a phosphatase that inhibits TCR signaling (81).

## Discussion

To our knowledge, this is the first study to correlate polygenic risk scores for any disease trait to T cell gene expression. We first calculated and validated a PRS for AD, comparing it to diagnostic data and pathological trait measurements. The AD PRS reached a maximum predictive value of 78.19%, within range of the predictive value of other AD PRSs reported in the literature (5–9). The predictive value of our AD PRS is likely inflated, however, due to the overlap of our target data cohort with the base data GWAS (34). We further validated the utility of our AD PRS by correlating it with neuropathological traits; showing that individuals with higher Braak scores, amyloid and tangle burden, and global pathology had, on average, higher polygenic risk for AD. Polygenic risk for AD has been shown elsewhere to correlate with neuropathological phenotypes such as higher amyloid burden as measured with PET (15,18), volume loss in brain regions such as the hippocampus and entorhinal cortex as seen on MRI (11,17), and levels of phosphorylated tau or amyloid beta in plasma or cerebrospinal fluid (19–21).

We then used the optimal *p*-value threshold from the AD PRS as a benchmark to correlate PRS for other immunological or disease traits to T cell gene expression from four sorted T cell subsets. Most PRS-associated genes were not shared widely across traits and cell types, and several autoimmune and psychiatric disease traits showed most genes in an inverse relationship with the PRS. Genes associated with the PRS for many traits in our dataset included genes related to T cell receptor signaling, T cell memory, and T cell activation. Correlating transcriptomic phenotypes across traits generally showed that traits within a particular category, such as autoimmune or psychiatric disease, had positively correlated gene expression patterns. Some exceptions, such as T cell PRS-associated gene expression in stroke being negatively correlated with that of other neurological disorders, but positively correlated with PRS-associated genes for psychiatric disease traits, could be interesting to follow up in replication studies, particularly using immune cells from stroke survivors. We know that the overall genetic architecture of several psychiatric disease traits features a notable degree of overlap (82), as do several subtypes of dementia (83). Our study suggests that PRS-associated T cell genes may reveal similar polygenic risk-mediated phenotypes across several related disease traits, though this should be confirmed in replication studies using relevant patient cohorts.

Pathway analysis mainly yielded biological processes related to cellular maintenance functions not specific to T cells. Several functionally intriguing pathways included genes known to affect migration and homing in T cells. In several disease states, T cell gene expression and clonality can change dramatically upon entry of T cells into target tissue. For example, in Parkinson’s and Lewy body dementia patients, T cells isolated from CSF presented a distinct transcriptomic signature from peripheral blood T cells, including upregulation of the chemokine receptor CXCR4 (3). Often, genetically-regulated gene expression changes in T cells are specific to particular T cell subsets (27,29–32,84) or activation states *in vitro* (29,30,85), suggesting that genotype-phenotype correlations specific to tissue microenvironment may exist as well. While our study profiles PRS-associated transcriptomic changes from peripheral T cells, future studies may optimize disease-relevant findings by isolating immune cells specifically from target tissue, such as the synovium in rheumatoid arthritis or post-mortem brain tissue samples from neurodegenerative disease decedents.

Polarization of PRS-associated genes by T cell subsets raises interesting questions. While most autoimmune disease traits had a majority of PRS-associated genes in CD4+ T cell subsets, with some psychiatric disease traits showing this trend in CD8+ T cell subsets, several traits showed differential enrichment of PRS-associated genes by cell type, depending on the direction of association between the gene and the PRS. Given that individuals with high polygenic risk for several autoimmune or psychiatric diseases showed widespread downregulation of T cell genes in our cohort (**Figure 1**), it would be interesting to determine whether this trend is replicated in T cells taken directly from patients with these conditions, and whether this phenomenon is specific to certain T cell subsets. Widespread downregulation of T cell effector genes, in particular, could represent a progression of T cells towards exhaustion in advanced disease.

The use of many contributing SNPs as input to the PRS often results in more generalized findings, compared to the single SNP-single gene approach of eQTL studies. This was certainly the case in our results, as only two PRS-associated genes remained significant after Bonferroni multiple testing correction at q = 0.05, unlike our trans-eQTL findings in the same cohort. Nevertheless, genes involved in modulation of T cell receptor signaling, cytokine signaling, T cell activation and memory, and T cell cytotoxicity arose in a few analyses. Studies that arise from other cohorts involving collection of genomic and gene expression data will be vital tools for comparing eQTL with polygenic risk-mediated transcriptomic phenotypes, especially those that collect gene expression data from specific tissues or cell types. Existing datasets from projects such as GTEx (86), which has already extensively profiled tissue-specific eQTL, could also be mined for polygenic risk-mediated gene expression changes in a variety of disease settings.

The importance of replication, in studies using genotype data alone or studies exploring genotype-phenotype correlation, cannot be overstated. The limitations of the present study certainly warrant future efforts in independent cohorts to confirm and expand on our stated findings. Specifically, while our subject pool for genotyping data was sufficient to establish well-powered findings for PRS calculation alone, the correlation with T cell gene expression was done in only 77 subjects. Also, as with our eQTL study, the use of CD45RO to distinguish naïve and memory T cells is not ideal. While we did not parse differences between CD45RO- and CD45RO+ populations to the same extent in the PRS study, analyses such as GSEA may have returned pathways that would not have been found with a sorting approach that properly distinguished memory from naïve cells.

The number of studies collecting genome sequencing or genotyping data alongside one or more quantitative traits will continue to expand, for AD and other disease cohorts. Thus far, studies using datasets like these tend to stratify PRSs into quantiles for comparison with quantitative traits, rather than using linear regression as we have done. In cohorts where well-powered comparisons are possible, it will be interesting to see whether quantitative phenotypes like gene expression or pathological biomarkers are better examined as a difference in means between stratified PRS quantile groups, or as a linear regression. The former approach would allow for a case-control type of study design, while the latter enables scalable predictions of how quantitative traits might vary with small differences in PRS. Either way, the utilization of PRSs in genotype-phenotype correlation studies will be invaluable for diseases where the contribution of T cells is coming to light, such as AD, and could be beneficial as well for diseases with sparsely established relevance to T cell biology, as with many psychiatric conditions.

In summary, our experiments show that T cell gene expression phenotypes can change with respect to polygenic risk for disease. We calculated a PRS for AD in our cohort and estimated its predictive value for disease, then calculated PRSs for 18 other diseases and traits. Correlating PRSs with gene expression, and determining PRS-associated genes with high overlap across traits and T cell subsets, detected genes involved in key aspects of T cell signaling and activation. We explored how PRS-associated T cell transcriptomic signatures compared between traits, and biological pathways and processes represented by PRS-associated genes. Finally, we found that several traits displayed differential polarization of their PRS-associated genes towards CD4+ or CD8+ T cell subsets, even when considering the direction of association between the PRS and gene expression. While we and others have previously detailed changes in T cell gene expression related to individual genetic variants, our analyses here show that many disease-associated variants can have an aggregate effect on T cell transcriptomic phenotypes as well.

## Materials and Methods

### The Rush Religious Orders (ROS) Study and Rush Memory and Aging Project (MAP)

ROS, started in 1994, enrolls Catholic priests, nuns, and brothers, without known dementia, aged 53 or older from more than 40 groups in 15 states across the USA. Since January 1994, more than 1,450 participants completed their baseline evaluation, of whom 87% are non-Hispanic white, and the follow-up rate of survivors and autopsy rate among the deceased both exceed 90%. MAP, started in 1997, enrolls men and women without known dementia aged 55 or older from retirement communities, senior and subsidized housing, and individual homes across northeastern Illinois. Since October 1997, more than 2,200 participants completed their baseline evaluation, of which 87% were non-Hispanic white. The follow-up rate of survivors exceeds 90% and the autopsy rate exceeds 80%. Both studies were approved by an Institutional Review Board of Rush University Medical Center. All participants signed informed consent for detailed annual clinical evaluation, an Anatomic Gift Act, and a repository consent to allow their data and biospecimens to be share. All ROSMAP participants agree to organ donation at death. A subset of 450 ROS and all MAP participants agreed to annual blood draw. Cryopreserved PBMC were stored at baseline and annually from 2008 to present. More detailed descriptions of ROS and MAP can be found in prior publications (25). ROSMAP resources can be requested at https://www.radc.rush.edu.

PBMC from 96 study autopsied participants were used in this study. 48 subjects were clinically and/or pathologically diagnosed with AD, while 48 subjects without dementia served as controls. Brain tissue from each subject was analyzed post-mortem to detect pathological signs of neuritic plaques and neurofibrillary tangles.

### Sample preparation, RNA-sequencing, and genotyping

Sample preparation and data acquisition for this study has been described previously (4). In brief, peripheral blood mononuclear cells (PBMCs) were isolated by Ficoll gradient centrifugation, then sorted by high-speed flow cytometry into the following T cell subsets: CD4+CD45RO-, CD4+CD45RO+, CD8+CD45RO-, and CD8+CD45RO+. Total RNA was extracted using buffer TCL (Qiagen), then RNA-seq libraries were prepared according to the Single Cell RNA Barcoding and Sequencing method originally developed for single cell RNA-seq (87), adapted for extracted total RNA. RNA was sequenced on the Illumina HiSeq using the Illumina HiSequsing the High-throughput 3’ Digital Gene Expression (DGE) library (87). Genes with maximum cell count value of at least three and non-zero values in over twenty percent of samples were included in differential expression analysis. Expression values were normalized to counts per million (CPM). The EdgeR package in R (88) was used to conduct differential expression, and Voom transformation (89) was applied to gene expression data. DNA for genotyping was extracted from whole blood or frozen post-mortem brain tissue and genotyped using the Affymetrix GeneChip 6.0 platform. Quality control of genotyping data was done with PLINK (90) (http://pngu.mgh.harvard.edu/~purcell/plink/), and imputation was done with MACH software (version 1.0.16a).

### Polygenic Risk Score (PRS) calculation

Summary statistics files from genome-wide association studies were used as the base data. Duplicate SNPs were removed from base data files, as were ambiguous SNPs for which the effect allele and other allele were complementary nucleotides (C with G or A with T). For summary statistics whose coordinates were found on genome build 38, LiftOver (http:/genome.ucsc.edu) was used to convert these coordinates to genome build 37, to match the target data. For traits with missing odds ratio values in the summary statistics, these were calculated from the beta values by using the exp() function in R. For traits with missing beta values, beta values were estimated using the sample size, Z-score, and allele frequencies using the equation below. The estimated beta values were then converted to odds ratio values using exp().

Genomic data from ROSMAP participants was used as the target data. These data were stored as three separate batches (ROSMAP_n1686, ROSMAP_n381, and ROSMAP_BU) due to separate genotyping batches, and initially processed individually by cohort. Quality control of target data was done with R and PLINK (90). First, SNPs were filtered to exclude those with a minor allele frequency (MAF) less than 0.01, a Hardy-Weinberg Equilibrium test p-value under 1 x 10^-6^, and SNPs missing in at least 1% of subjects. Individuals missing over 1% of SNPs in their genotyping data were also excluded at this stage. We then pruned highly correlated SNPs using a window size of 200 variants, a step size of 50 variants at a time, and filtering out any SNPs with an LD *r*^2^ value above 0.25. Subjects with heterozygosity F coefficients greater than three standard deviations from the mean, with differences between reported sex and sex chromosomes, or with a first or second degree relative in the sample, were excluded.

After these quality control steps on individual batches, we used PLINK to merge ROSMAP_n1686, ROSMAP_n381, and 15 subjects from ROSMAP_BU. We limited the inclusion of subjects from ROSMAP_BU to subjects with T cell gene expression data whose AD PRS was not an outlier in the overall distribution. This was because of the high number of unique SNPs in the ROSMAP_BU cohort relative to ROSMAP_n1686 and ROSMAP_n381, which we suspect is due to low-quality imputation of rare variants. In the merged dataset, we excluded SNPs with MAF < 0.01 and SNPs missing in at least 5% of subjects. The merged dataset was then used as target data in further PRS calculation.

We used PRSice-2 (33) for PRS calculation. PRSice-2 performs clumping of input variants using parameters of a 250 kb window, p-value threshold of 1, and *r*2 value threshold of 0.1. The software also automatically performs strand-flipping for SNPs whose alleles mismatch between base and target data. Individual SNPs were weighted by odds ratio for PRS calculation, and age and sex were used as covariates. For the AD PRS, clinical AD diagnosis was used as the input phenotype, and SNPs within 1 Mb of the APOE locus were excluded as input in PRS calculation. We also input a range of P-value thresholds for SNP inclusion from 5 x 10^-8^ (genome-wide significant SNPs only) to 1 (all SNPs), yielding a set of PRS scores as output for each individual.

### Validation of AD PRS against clinical and pathological data

The pROC package (91) was used to calculate receiver operator characteristic (ROC) curves for the AD PRS against clinical AD diagnosis and against pathological AD diagnosis, at each SNP significance threshold. We also used this package to calculate the predictive value of the AD PRS from the area under the ROC curves, to determine which significance threshold yielded the highest predictive value. We compared the distribution of AD PRS scores at this significance threshold between AD and non-AD subjects using student’s T test. We also ran comparisons of the AD PRS at this significance threshold against Braak score using nonparametric one-way ANOVA, and quantitative pathological measurements (amyloid plaque burden, tau tangle burden, and global pathology measurement) using linear regression with age, sex, clinical AD status, pathological AD status, and the first ten principal components from genotyping data as covariates.

### Detection and pathway analysis of PRS-associated genes

Genotyping data and T cell RNA-sequencing data were available for 78 subjects, with one subject excluded whose AD PRS was an outlier. For each trait and each T cell subtype, we used linear regression of gene expression counts against PRS scores to detect PRS-associated genes. Genes expressed in fewer than 20% of subjects, or with a maximum expression count under 3, were excluded from PRS association analyses. Because non-AD traits did not have relevant phenotypic data for PRS validation, we used the same significance threshold (p = 1) as the optimum PRS for all traits. For pathway analysis of PRS-associated genes, we used Gene Set Enrichment Analysis (GSEA, (59)) separately for each T cell subtype and each trait. GSEA was run using the t-value from the PRS-gene association as the ranking metric, using the default value of 1,000 permutations and restricting the gene sets to those with 15-200 genes, using the GO Biological Processes 2022 database (60). We then detected biological pathways which were significantly over-represented or under-represented at a significant level of 0.05, after correcting for false discovery rate.

## Authors Contributions

W.E. conceptualized and implemented the immunological study design. D.A.B. contributed the human blood samples and clinical and neuropathological data and critically reviewed the paper. D.D., W.E., and B.V., performed statistical analyses, interpretation of results, and wrote the manuscript. S.T and L.Y. provided technical guidance to optimize PRS calculations across traits. S.T, L.Y, J.S and D.A.B edited and revised the manuscript. All authors read and approved the final manuscript.

## Acknowledgments

We thank the participants in the Religious Orders Study and the Rush Memory and Aging Project for donating their data and biospecimen. This work was supported by the US National Institutes of Health grants R01AG067581 and R01NS124771 and the Ludwig Family Foundation (to W.E), and P30AG10161, P30AG72975, R01AG15819, R01AG17917, and U01AG61356 to D.A.B.. D.D. is a recipient of a TL1 Precision Medicine Predoctoral Fellowship (TL1TR001875).

## Abbreviations

Aβ: Amyloid beta
AD: Alzheimer’s disease
ALS: Amyotrophic lateral sclerosis
ANOVA: Analysis of variance
clinAD: Clinical diagnosis of Alzheimer’s disease
CNS: Central nervous system
CPM: Counts per million
CSF: Cerebrospinal fluid
DGE: Digital gene expression
DNA: Deoxyribonucleic acid
eQTL: Expression quantitative trait loci
GO: Gene ontology
Gpath: Global pathology score
GSEA: Gene set enrichment analysis
GWAS: Genome-wide association study
LD: Linkage disequilibrium
MAF: Minor allele frequency
Mb: Megabase
MRI: Magnetic resonance imaging
MS: Multiple sclerosis
pathoAD: Pathological diagnosis of Alzheimer’s disease
PBMC: Peripheral blood mononuclear cell
PET: Positron emission tomography
PD: Parkinson’s disease
PRS: Polygenic risk score
PTSD: Post-traumatic stress disorder
RNA: Ribonucleic acid
ROC: Receiver operating characteristic
ROSMAP: Religious Orders Study/Memory and Aging Project
SNP: Single nucleotide polymorphism
TCR: T cell receptor

## Legends to Supplementary Figures and Tables

**Supplementary figure 1. Validation of the AD PRS against diagnostic data and pathological trait measurements. A)** ROC curves, colored by the SNP p-value threshold used to calculate the PRS, showing the predictive value of each set of PRSs for AD against clinical diagnosis of AD. **B)** ROC curves showing the predictive value of the AD PRS against pathological diagnosis of AD. **C)** Comparison of the PRS score distribution by Braak score. **D-F)** Scatter plot of the AD PRS against quantification of amyloid burden, tau tangle score, and global pathology score (gpath).

**Supplementary figure 2. Summary of the overlap among PRS-associated genes across traits and T cell subsets. A)** Histogram showing the number of genes (y axis) found associated with the PRS for a given number of traits (x axis) across T cell subsets. **B)** Stacked bar chart showing the number of genes (y axis) found associated with the PRS for a given number of traits (x axis) within T cell subsets, with cell type given by color.

**Supplementary table 1.** All PRS-associated genes, with associated effect sizes, unadjusted *p*-values, and adjusted *p*-values after multiple testing correction by Benjamini-Hochberg false discovery rate or Bonferroni correction. Each tab is for one trait and one T cell subset (C1 is CD4+CD45RO-, C2 is CD4+CD45RO+, C3 is CD8+CD45RO-, C4 is CD8+CD45RO+).

**Supplementary table 2.** Quantities of Pearson’s r values for figure 2 are shown above the yellow diagonal. NA values indicate correlations that were insignificant after Bonferroni multiple testing correction. Absolute numbers of shared significant PRS-associated genes between any two traits are shown below the yellow diagonal.

**Supplementary table 3.** All GSEA pathways from all traits and T cell subsets that are significantly over- or under-represented among PRS-associated genes after multiple testing correction. SIZE refers to the number of genes in the pathway, t.value.sign refers to the direction of association of pathway genes with the PRS for the given trait.

**Supplementary table 4.** Quantities used to obtain the figure 4 heatmap, including absolute counts of PRS-associated genes in CD4+ and CD8+ subtypes for each trait, percentage of total, and percent differences between CD4+ and CD8+ subtypes.

