## Supplementary figures and images for "Polygenic Risk Associated with Alzheimer’s Disease and Other Traits Influences Genes Involved in T Cell Signaling and Activation"

### Supplementary figure 1

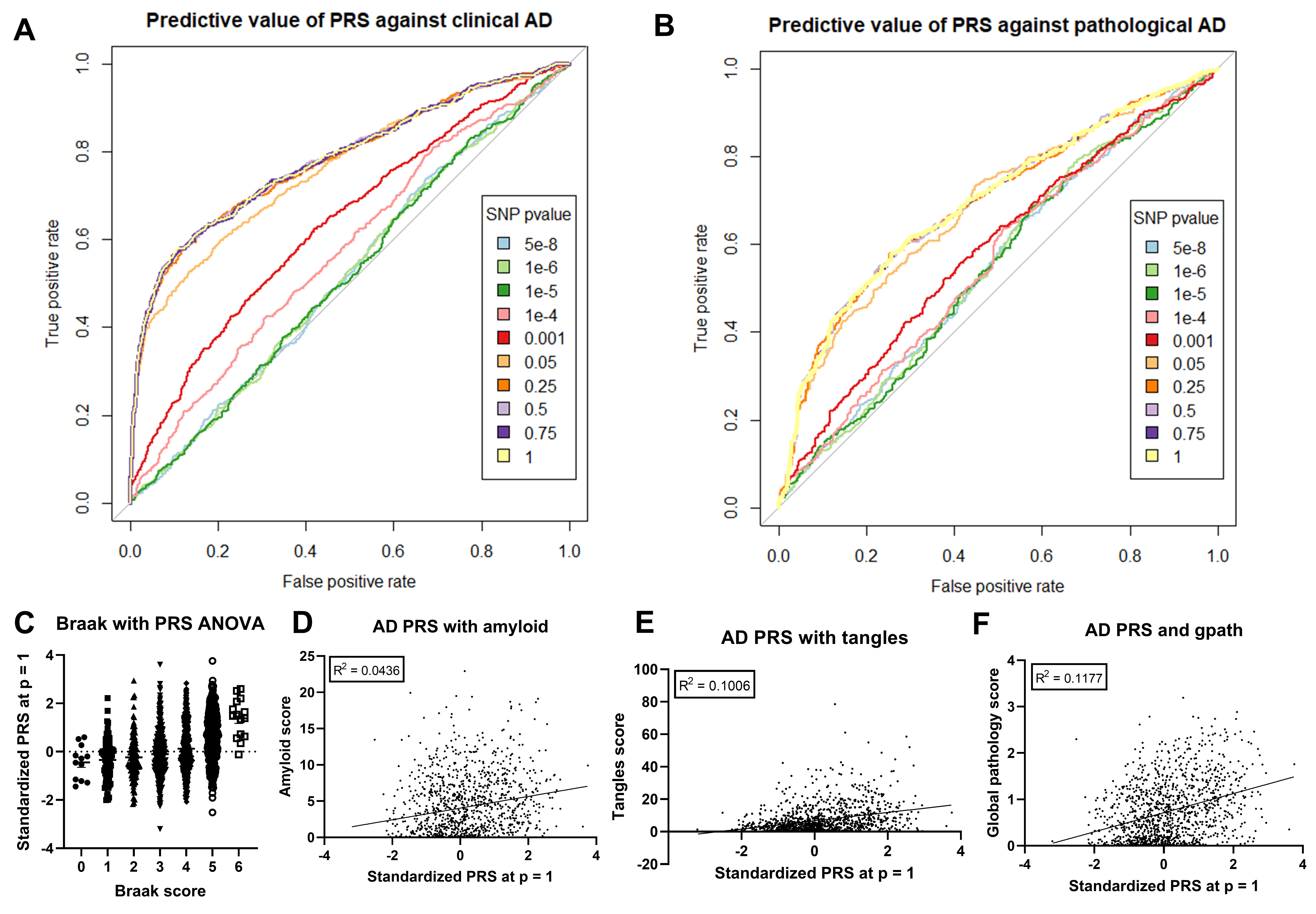

### Supplementary figure 2

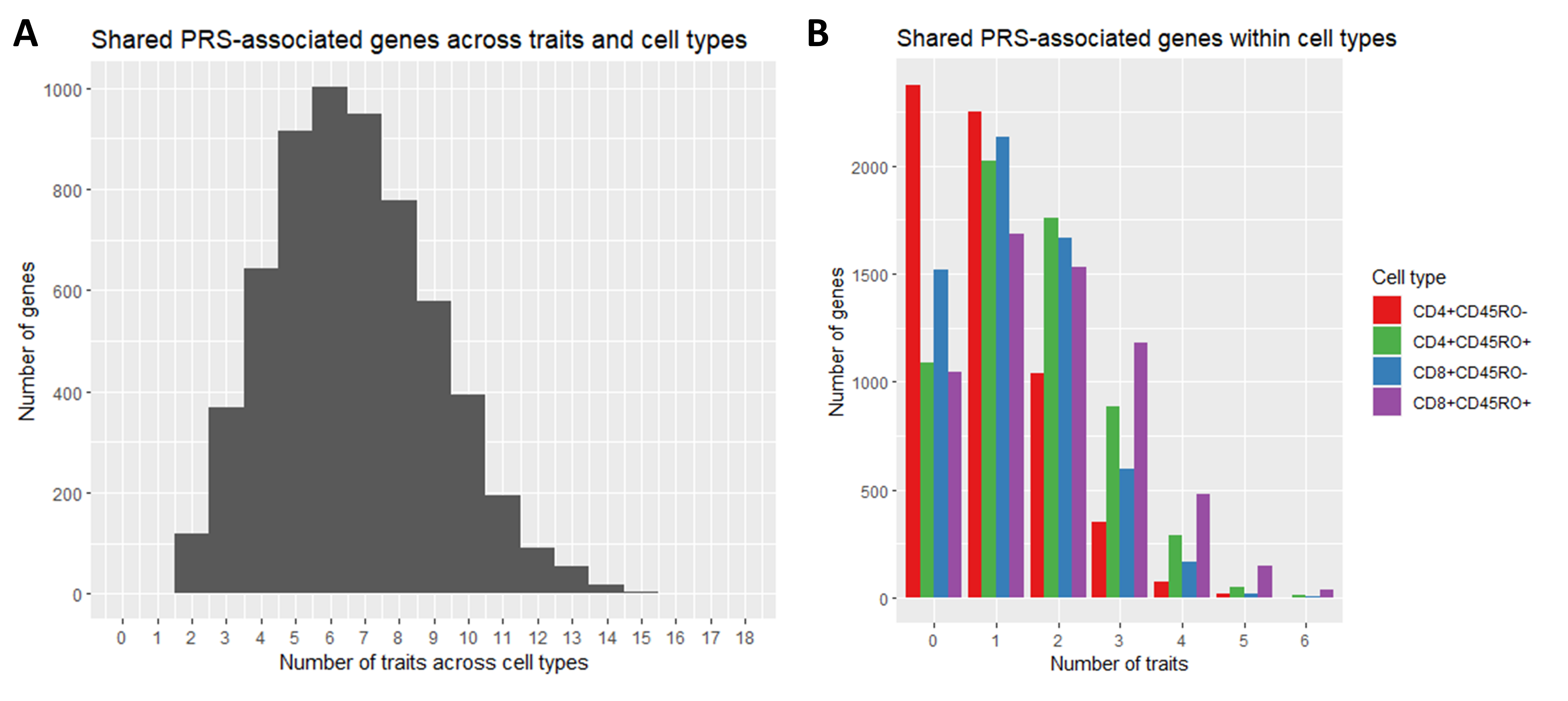
